# Enhancing working memories using repetitive TMS

**DOI:** 10.64898/2026.10.22.761198

**Authors:** Mattia F. Pagnotta, Mark D’Esposito

**Affiliations:** Helen Wills Neuroscience Institute, University of California Berkeley, Berkeley, CA, USA; Department of Psychology, University of California Berkeley, Berkeley, CA, USA

## Abstract

Working memory, the ability to retain and manipulate information no longer present in the environment, is fundamental for goal-directed behavior (Baddeley, 2012; Baddeley & Hitch, 1974; D’Esposito, 2007; D’Esposito & Postle, 2015). Alpha oscillations (∼8–12 Hz) have two apparently contradictory roles in working memory. Alpha activity gates irrelevant information through inhibition, a role causally supported by rhythmic transcranial magnetic stimulation (TMS) of parietal cortex at alpha frequency when a retrospective cue renders a memory item irrelevant (Bonnefond & Jensen, 2012; Riddle et al., 2020). Yet alpha activity also tracks the content of memory and binds the features of memorized objects during maintenance (Foster et al., 2016; Pagnotta et al., 2026). Whether alpha oscillations causally support the fidelity of memory representations during maintenance is untested. Here, 24 participants received short trains of repetitive TMS (rTMS) at theta (6 Hz) or alpha frequency (10 Hz), or in an arrhythmic control pattern, during the second half of the delay of a color working memory task, over the left posterior inferior frontal sulcus or the left intraparietal sulcus. A mixture model separated memory precision from swap errors and guesses. Alpha stimulation of the intraparietal sulcus specifically increased memory precision, whereas rhythmic stimulation of either site reduced response error, guesses and swaps irrespective of frequency. These results provide causal evidence that alpha oscillations support the fidelity of working memory representations during maintenance, and show that rhythmic stimulation at the appropriate frequency, site and time can enhance human working memory.

## RESULTS AND DISCUSSION

Alpha oscillations (∼8–12 Hz) have been assigned two roles in working memory that are, at first sight, at odds. Causal evidence from rhythmic transcranial magnetic stimulation (TMS) locked to a retrospective cue showed that parietal alpha gates the content of working memory by suppressing information that is no longer relevant (Riddle et al., 2020). This is consistent with the view that alpha provides functional inhibition (Bonnefond & Jensen, 2012; Klimesch et al., 1999). Yet the topography of alpha power tracks the specific content held in memory throughout the delay (Bender et al., 2025; Foster et al., 2016), and alpha phase coding binds the features of memorized objects, with binding errors arising from unstable phase-locked activity (Pagnotta et al., 2026), as predicted by attractor models in which oscillatory synchrony organizes competing memories (Barbosa et al., 2021). Whether alpha oscillations causally support the fidelity of working memory representations during maintenance has not been tested. We tested this hypothesis by delivering short trains of rhythmic TMS at theta (6 Hz) or alpha frequency (10 Hz), or an arrhythmic control train, during the second half of a 2 s memory delay, over the left posterior inferior frontal sulcus (pIFS) or the left intraparietal sulcus (IPS), in 24 participants (Figure 1; see Methods). Participants reported the color of a cued item six times in each trial (Jabar et al., 2023), and a mixture model (Bays et al., 2009; Schneegans & Bays, 2016) separated memory precision (the concentration parameter κ) from swap errors and guesses. All measures were expressed relative to arrhythmic stimulation at the same site and in the same session, to isolate frequency-specific effects from non-specific effects of stimulation (Riddle et al., 2020).

**Figure 1.**
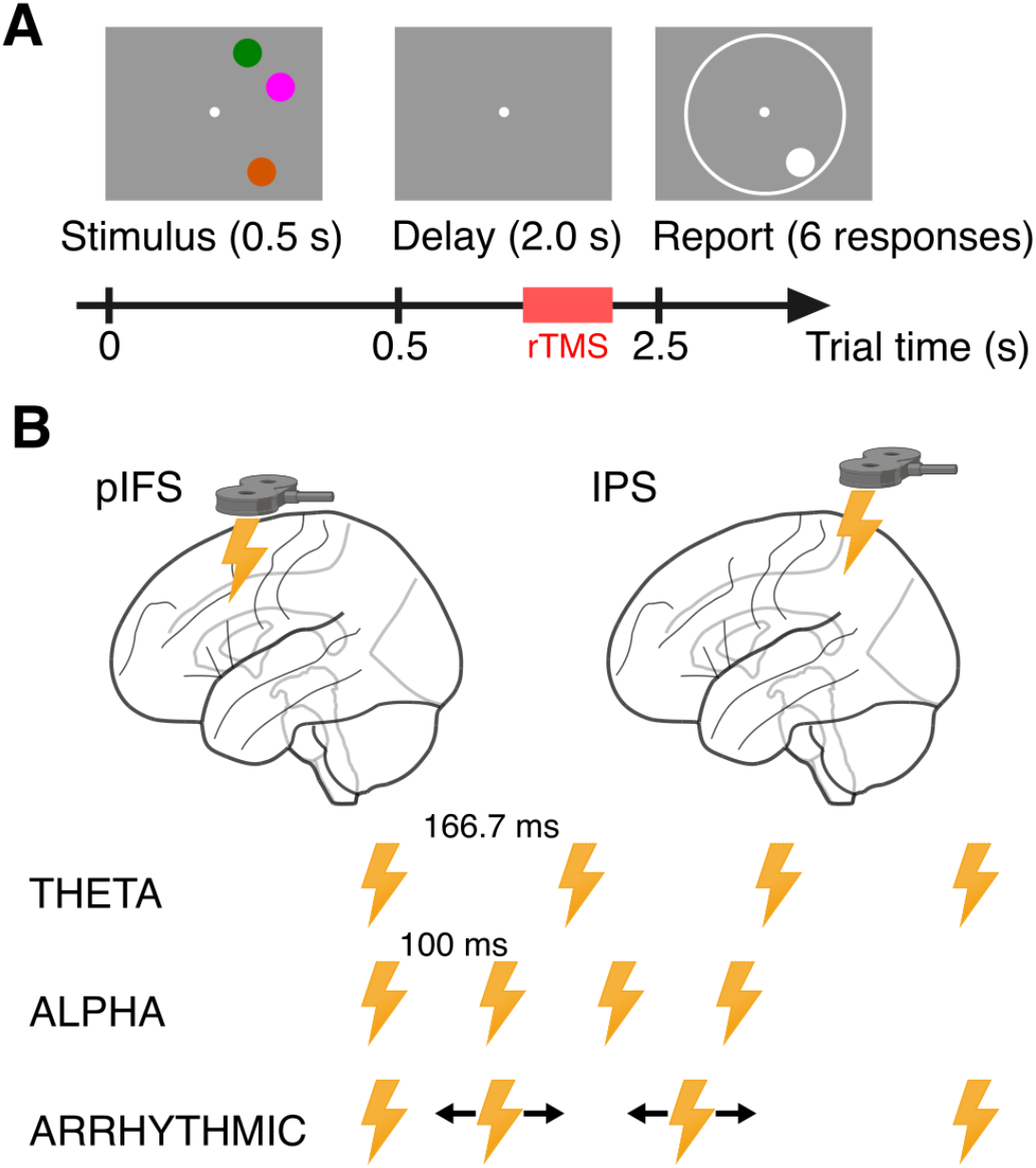
Working memory task and repetitive TMS (rTMS) protocol. (A)Trial sequence. Each trial began with a memory array of three colored circles presented for 0.5 s in one visual hemifield (left or right, equiprobable) around a central fixation dot, followed by a 2 s delay during which only the fixation dot remained on screen. At the end of the delay, a circle appeared at the location of one of the three items, cueing it as the target, together with a color wheel (schematized here as a white ring; the actual wheel displayed 360 isoluminant colors, randomly rotated on every trial). Participants reported the color of the cued item by moving the mouse along the wheel, which filled the circle with the color under the cursor, and confirmed with a click; six consecutive responses were collected on each trial (self-paced). Repetitive transcranial magnetic stimulation (rTMS; red band) was delivered during the second half of the delay: trains began 1.0–1.2 s after the offset of the memory array and ended no later than 1.7 s, i.e., at least 0.3 s before the report screen. Pre-stimulus fixation (1 s), feedback (1 s), and inter-trial interval (0.5 s) are omitted; the timeline is not to scale. (B)Stimulation sites and pulse patterns: Top: rTMS was applied over the left posterior inferior frontal sulcus (pIFS; MNI -46, 0, 38) or the left intraparietal sulcus (IPS; MNI -38, -50, 44), one site per session, with the order of sites counterbalanced across participants. Bottom: on every trial a train of four pulses (lightning bolts) at 110% of the individual resting motor threshold was delivered in one of three patterns, intermixed across trials. Theta: 6 Hz (inter-pulse interval of 166.7 ms; train duration of 500 ms). Alpha: 10 Hz (inter-pulse interval of 100 ms; train duration of 300 ms). Arrhythmic: first and last pulses 500 ms apart, with the two intermediate pulses at random times on each trial (double-headed arrows; all inter-pulse intervals between 20 and 400 ms), serving as a control for non-frequency-specific effects of stimulation at each site.

### Alpha-frequency stimulation of parietal cortex during maintenance enhances memory precision

Rhythmic stimulation changed memory precision in a manner that depended on where and at which frequency it was delivered (Figure 2A). A three-way repeated-measures ANOVA on the change in concentration parameter κ (rhythmic minus arrhythmic), with factors stimulation frequency (theta, alpha), stimulation site (pIFS, IPS), and presentation hemifield (left, right), revealed a main effect of site (F(1,23)=7.915, p=0.0098; d=0.560): stimulation of IPS increased precision relative to arrhythmic stimulation, whereas stimulation of pIFS did not. This effect was qualified by a frequency-by-site interaction (F(1,23)=4.809, p=0.0386; d=0.172). Post hoc comparisons (Tukey–Kramer) showed that alpha stimulation of IPS improved precision relative to alpha stimulation of pIFS (p=0.0032; d=0.662), and relative to theta stimulation of IPS (p=0.0498; d=0.233), although the latter difference was small and at the threshold of significance. Accordingly, the distribution of responses around the target color was narrowest after alpha stimulation of IPS and broadest after alpha stimulation of pIFS (Figure 2A, right). Precision did not depend on the hemifield of the memory array, and no other main effect or interaction reached significance. Thus, the benefit of alpha stimulation was specific to the parietal site and was not mirrored by theta stimulation of the same site.

**Figure 2.**
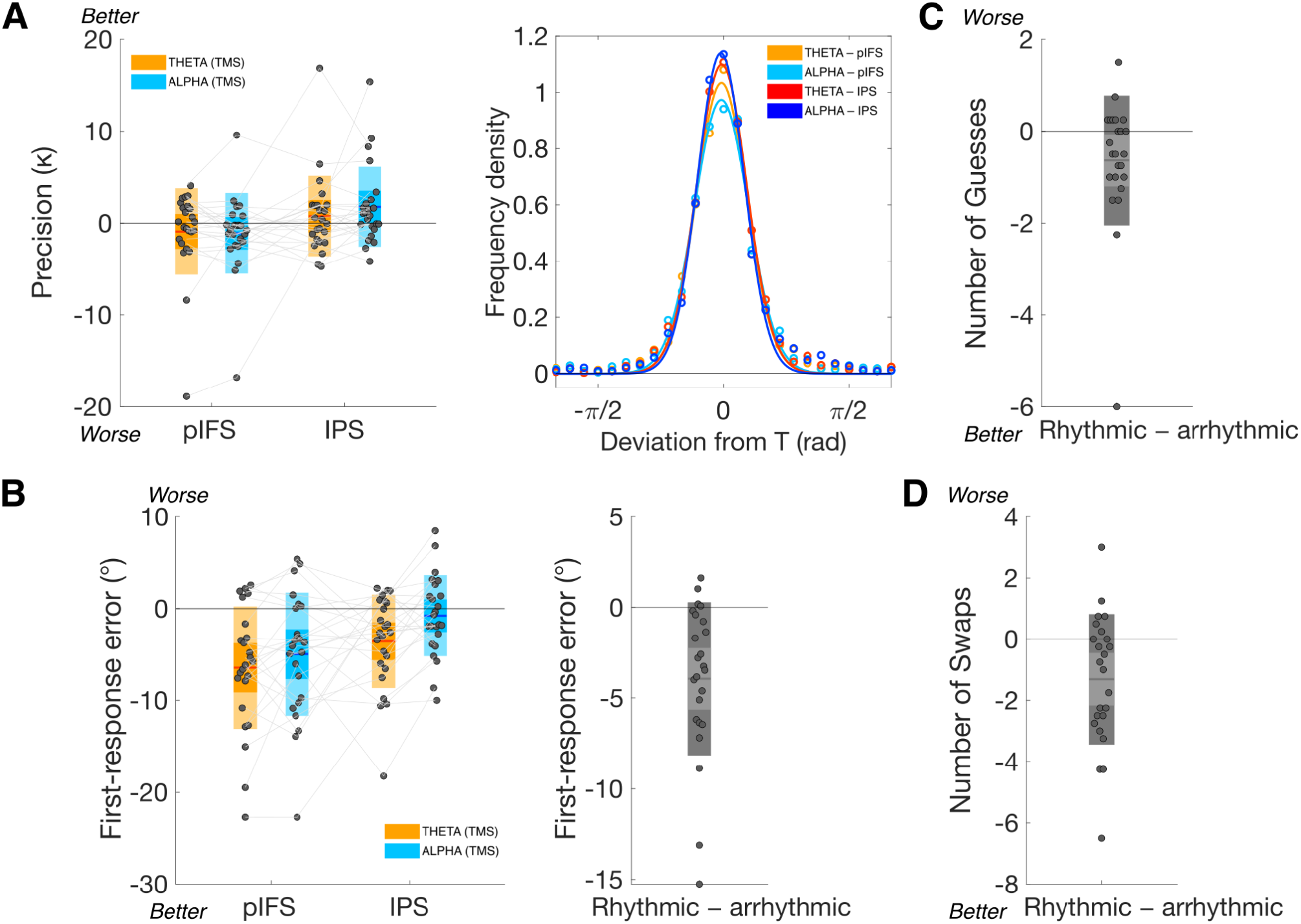
Rhythmic TMS during working memory maintenance enhances precision in a site- and frequency-specific manner and reduces errors. (A)Memory precision. Left: change in the concentration parameter κ of the von Mises component of the mixture model, computed for each participant as the difference between each rhythmic condition and the arrhythmic condition from the same session (rhythmic–arrhythmic), for theta (orange) and alpha (cyan) stimulation of pIFS and IPS. Left- and right-hemifield trials were pooled. Dots represent individual participants (N=24), jittered horizontally; grey lines connect the values of the same participant across conditions; the colored horizontal line is the group mean, the darker box its 95% confidence interval (mean ± 1.96 SEM), and the lighter box ± 1 SD. The black line at zero marks no change relative to arrhythmic stimulation; “Better” and “Worse” indicate the direction of improved performance. Right: distribution of the deviation of responses from the target color (all six responses per trial, hemifield pooled) for the four rhythmic conditions. Circles are the mean probability density across participants of 10° bins; lines are the von Mises-shaped curves fitted to the group means for display only. (B)First-response error. Left: change in the mean absolute error of the first response on each trial, in degrees of color space, relative to the arrhythmic stimulation; conventions as in panel (A). Right: for each participant, the mean of the four rhythmic–arrhythmic differences shown on the left, i.e., the overall effect of rhythmic stimulation irrespective of frequency and site; the grey line represents the mean, boxes the 95% confidence interval and ± 1 SD. (C)Number of guess trials and (D) number of swap trials, as classified by the mixture model, expressed as in the right panel of (B): the per-participant mean change relative to arrhythmic stimulation across the four rhythmic conditions, with negative values indicating fewer guesses or swaps. Statistics are reported in the text. *pIFS*, posterior inferior frontal sulcus; *IPS*, intraparietal sulcus; *SEM*, standard error of the mean; *SD*, standard deviation.

### Rhythmic stimulation reduces response error, guesses and swaps irrespective of frequency and site

The absolute error of the first response on each trial showed a different pattern (Figure 2B). The intercept of the ANOVA was significant (F(1,23)=20.935, p=0.0001), indicating that first-response error was lower after rhythmic than arrhythmic stimulation irrespective of frequency, site and hemifield (Figure 2B, right). The size of this reduction depended on frequency (main effect, F(1,23)=7.611, p=0.0111; d=0.463), being larger after theta than after alpha stimulation, and on site (main effect, F(1,23)=7.185, p=0.0133; d=0.666), being larger after stimulation of pIFS than of IPS. The two factors did not interact, and the hemifield had no significant effect. The pattern of first-response error was thus nearly the reverse of that for precision: the conditions that most reduced the error of the first report (theta, pIFS) were not those that most increased the concentration of responses around the target color (alpha, IPS).

The numbers of guess trials and swap trials, as classified by the mixture model, were likewise reduced by rhythmic relative to arrhythmic stimulation (guesses: intercept, F(1,23)=4.898, p=0.0370; swaps: intercept, F(1,23)=9.162, p=0.0059; Figure 2C–D). Neither measure depended on stimulation frequency, stimulation site or presentation hemifield, and no interaction reached significance.

Two aspects of these results dissociate. Memory precision was enhanced in a site- and frequency-specific manner, by alpha stimulation of the parietal cortex. First-response error, guesses and swaps, in contrast, showed a general benefit of rhythmic over arrhythmic stimulation, with the error reduction tilted toward theta stimulation and toward the frontal site. Because first-response error pools accurate reports, swaps and guesses, part of its reduction reflects the lower incidence of gross errors. The concentration parameter κ, in contrast, indexes the spread of responses attributed to the target component once swaps and guesses are accounted for. The specific improvement in precision after alpha stimulation of IPS cannot be explained by fewer swaps or guesses, since these were reduced irrespective of frequency and site.

### Alpha oscillations support the fidelity of working memory representations during maintenance

These findings provide causal evidence that alpha oscillations in the parietal cortex support the fidelity of working memory representations during maintenance. In our design, stimulation was delivered after encoding and before the cue, with no distractor and with all memory items in the same hemifield, so that there was no irrelevant information to be gated when the stimulation trains arrived. Under these conditions, alpha-frequency stimulation of IPS increased the precision with which the cued item was later reported. This result is the causal counterpart of the correlational observations that parietal-occipital alpha activity carries the content of working memory throughout the delay (Bender et al., 2025; Foster et al., 2016), and it complements the evidence that alpha phase coding binds object features during maintenance, with binding errors emerging from transient phase instabilities (Pagnotta et al., 2026). In the attractor model that motivated that work, oscillatory phase relationships organize competing memories within and across feature-selective networks (Barbosa et al., 2021). An externally imposed alpha rhythm may stabilize these relationships, which would be expected to sharpen the stored representations rather than change what is stored. Precision is the measure that a phase-coding account predicts alpha to affect, and it is the measure that alpha stimulation selectively improved. These results extend the causal role of alpha oscillations, previously established for the control of working memory at the time of a retrocue (Riddle et al., 2020), to the maintenance of its content.

An alternative account holds that increases in alpha activity during the delay reflect functional inhibition that protects maintained representations from interference (Bonnefond & Jensen, 2012; Klimesch et al., 1999; Wianda & Ross, 2019), a view that remains the most common interpretation of delay-period alpha (Pavlov & Kotchoubey, 2022). In Riddle et al. (Riddle et al., 2020), alpha stimulation of IPS benefited performance by suppressing the representation that the retrocue had rendered irrelevant. A pure suppression account would not predict a benefit of alpha stimulation when no external input or irrelevant representation competes with the cued item, as was the case in the present study. Our data favor a contribution of alpha to the maintained representation itself. We note, however, that the two accounts need not be mutually exclusive. If the inhibition implemented by alpha is the lateral inhibition between competing representations from which phase coding emerges (Barbosa et al., 2021), then protecting a representation from internal interference and stabilizing its phase code may be two descriptions of the same mechanism. Our behavioral data cannot distinguish between them, and doing so will require concurrent neurophysiological recordings.

The specificity of the precision effect to the parietal site is consistent with the distributed organization of working memory, in which sensory and parietal regions retain stimulus information while prefrontal cortex biases and selects among representations (Christophel et al., 2017; D’Esposito, 2007; D’Esposito & Postle, 2015). Perturbation of retinotopic parietal cortex during the delay degrades the fidelity of the remembered location, whereas perturbation of frontal cortex degrades the planned response (Mackey & Curtis, 2017), and perturbation of early visual cortex during maintenance degrades memory in a retinotopically specific manner (Dake & Curtis, 2025). Our results add that the fidelity of a parietal representation can be enhanced, and not only disrupted, when stimulation is delivered at alpha frequency. Stimulation of pIFS did not improve precision at either frequency. This frontal site, at the junction of the inferior frontal and precentral sulci, differs from the anterior middle frontal gyrus targeted by Riddle and colleagues (Riddle et al., 2020), and lateral prefrontal cortex is better characterized as controlling the use of working memory contents than as storing them (Kiyonaga et al., 2024).

Rhythmic stimulation also reduced first-response error, guesses and swaps relative to arrhythmic stimulation, irrespective of frequency and site. Because all measures were difference scores, a benefit of rhythmic stimulation and a cost of arrhythmic stimulation are indistinguishable in our design, as in the design of Riddle et al. (Riddle et al., 2020). Online rTMS most often impairs rather than improves performance (Beynel et al., 2019), and arrhythmic trains may be the more disruptive pattern, for example because their timing cannot be anticipated. Rhythmic trains, in contrast, are temporally predictable and the alignment of ongoing oscillations to predictable events is itself associated with better performance (Bonnefond & Jensen, 2012). The larger reduction of first-response error after theta stimulation and after stimulation of pIFS is compatible with a role of frontal theta in the prioritization and readout of working memory contents (Riddle et al., 2020; Wallis et al., 2015), given that the first report is the most immediate translation of the memory into action. We refrain from a stronger interpretation, since theta oscillations are difficult to detect in human scalp recordings once aperiodic neural activity is accounted for (van Engen et al., 2026), and since our frontal target differed from that of previous work.

Several limitations qualify these conclusions. We did not record neural activity during stimulation, so we cannot establish whether 10 Hz trains entrained endogenous alpha oscillations, nor whether the behavioral effect reflects oscillatory rather than aperiodic activity, a distinction that recent work has shown to matter (Bender et al., 2025; Preston et al., 2025; van Engen et al., 2026). Stimulation was restricted to the left hemisphere, and its effect did not depend on the hemifield of the memory array; the benefit of parietal alpha stimulation therefore appears to operate at the level of a bilateral network rather than retinotopically, in contrast with the contralateral effects of disruptive stimulation (Dake & Curtis, 2025; Mackey & Curtis, 2017). We used a single intensity and a single window late in the delay, and the advantage of alpha over theta stimulation at IPS was small. Future work combining rhythmic TMS with electroencephalography, varying the timing of the stimulation within the delay, and targeting right hemisphere and sensory sites will be necessary to establish the oscillatory mechanism and its spatial extent.

In summary, alpha oscillations appear to have two lives in working memory: a gating role at the time of selection, and a role in sustaining the fidelity of representations during maintenance. By delivering rhythmic stimulation at the right frequency, site and moment, we enhanced rather than disrupted human working memory, an outcome that remains uncommon for online rTMS (Beynel et al., 2019) and that points to the maintenance period as a window for neuromodulation of memory.

## METHODS

### EXPERIMENTAL MODEL AND STUDY PARTICIPANT DETAILS

Twenty-seven healthy adults were recruited from the University of California, Berkeley community. The study was approved by the Committee for Protection of Human Subjects at the University of California, Berkeley, and all participants gave written informed consent before taking part. Participants were compensated at a rate of $20 per hour, independent of task performance. Before enrollment, all candidates were screened for contraindications to magnetic resonance imaging (MRI) and transcranial magnetic stimulation (TMS) (Rossi et al., 2009). Exclusion criteria for MRI included metal implants or fragments anywhere in the body (including surgical hardware, pacemakers, stents, or metal introduced through metalwork or injury), non-removable hearing aids or cochlear implants, non-removable metal dental work, a history of claustrophobia, and any prior exclusion from MRI. Exclusion criteria for TMS included a personal or family history of epilepsy, seizures, or convulsions; a history of fainting, stroke, serious head injury, brain surgery, or other neurological disorder; metal in the head or skull; implanted medical devices; treated hypertension; hearing problems or tinnitus; a serious medical condition; use of medication for a psychological condition within the previous three months; chronic or recent increases in stress, anxiety, or sleep disturbance; and drug or alcohol use in the preceding 24 hours. Participants were additionally screened for color blindness and reported normal or corrected-to-normal vision.

Three participants were excluded after enrollment: two because they were too sensitive to the stimulation to complete the experimental sessions, and one because a reliable motor threshold could not be established. The final sample comprised 24 participants (age range 18–37 years, *M*=22.42, *SD*=4.07; 14 female and 10 male; 19 right-handed, 4 left-handed, 1 ambidextrous). The sample size was determined a priori based on previous rhythmic-TMS studies with comparable within-subject designs (Kiyonaga et al., 2024; Riddle et al., 2020).

## METHOD DETAILS

### Stimuli, design, and procedures

Each participant completed three sessions on separate days. In the first session we determined the individual resting motor threshold (see Non-invasive brain stimulation below); in the second and third sessions participants performed the working memory task while receiving online repetitive TMS (rTMS) to either the left posterior inferior frontal sulcus (pIFS) or the left intraparietal sulcus (IPS), one site per session, with the order of sites counterbalanced across participants (12 participants received pIFS stimulation first). The two rTMS sessions were approximately one week apart. A high-resolution anatomical MRI scan, acquired before the first session or retrieved from a previous study on campus with the participant’s permission, was used for neuronavigation in all three sessions.

The working memory task (Figure 1A) was adapted from our previous study (Pagnotta et al., 2026), with the cue and report features reversed: participants were cued by an item’s location and reported its color. Stimuli were presented with Psychtoolbox-3 (Kleiner et al., 2007) in MATLAB on a V176L LCD Monitor (Acer Inc., Taipei, Taiwan; 17 inches, 5:4 square aspect ratio, resolution 1280x1024 SXGA at 60 Hz) viewed from 78–82 cm, against a grey background. A chin rest was used to stabilize the participant’s head and maintain a fixed viewing distance. Each trial began with a central fixation dot (0.2° of visual angle, DVA, in radius) presented for 1 s. The memory array, three colored circles (0.5 DVA radius) centered 4.5 DVA from fixation, was then presented for 0.5 s, followed by a 2 s delay during which only the fixation dot remained on screen. On each trial all three items appeared within the same hemifield (left or right, equiprobable), at integer polar angles at least 15° away from the vertical meridian and at least 20° apart from one another, to prevent physical overlap and to avoid swap errors driven by misperception of location. Colors were drawn without replacement from a 360-step color wheel defined as an isoluminant circle in CIELAB space (L* = 54), with a minimum separation of 20 steps between items to avoid swaps driven by color misperception.

At the end of the delay, a grey circle appeared at the location of one of the three items, designating it as the target, together with a color wheel (annulus with inner radius of 5.5 DVA and outer radius of 6.5 DVA) whose orientation was randomly rotated on every trial. Participants moved the mouse along the wheel, and the circle at the cued location was continuously filled with the color under the cursor; a click confirmed the response. Rather than a single response, participants provided six responses on each trial, following a wagering procedure that allows the precision of a memory to be estimated from the spread of repeated reports (Jabar et al., 2023). Each confirmed response added a Gaussian-shaped “bet” (σ = 3 wheel steps) to a cumulative white profile drawn around the outside of the wheel, so that participants could distribute their bets across the colors they considered plausible. The first response was weighted twice as heavily as the remaining five, and placing further bets on an already-wagered color yielded diminishing returns. The cursor was repositioned to a random point on the wheel before each response. Points earned on each trial were proportional to the height of the cumulative profile at the true color of the target (height^0.93 × 100), and were shown together with the running total during a 1 s feedback display that also showed the target’s true color. Trials were separated by a 0.5 s inter-trial interval. The report period was self-paced. Points served only to motivate accurate and well-calibrated responding and were not converted into monetary reward.

Each rTMS session consisted of 8 blocks of 30 trials (240 trials in total), with self-paced breaks between blocks. The six conditions defined by presentation hemifield (left, right) and rTMS pattern (theta, alpha, arrhythmic; see below) were balanced within each block (5 trials per condition per block, 40 per condition per session) and presented in random order. Before the first rTMS session participants were familiarized with the task using a practice block consisting of 6 trials, without stimulation.

### MRI data and selection of targets

MRI data were collected at the Henry H. Wheeler, Jr. Brain Imaging Center at the University of California, Berkeley, on a Siemens 3T MAGNETOM Trio (Erlangen, Germany) with a 32-channel receive-only head coil. A whole-head T1-weighted anatomical image was acquired with a magnetization-prepared rapid gradient-echo (MPRAGE) sequence (160 sagittal slices, 1 mm isotropic voxels, TR = 2.3 s, TE = 2.98 ms, flip angle = 9°, anterior-to-posterior phase encoding, GRAPPA acceleration factor 2).

The two cortical targets for rTMS were defined from a meta-analysis of 1,091 neuroimaging studies associated with the term “working memory” in the Neurosynth database (association-test z-map, false discovery rate corrected at q = 0.01) (Yarkoni et al., 2011). We selected the peak voxel (maximum z) of the left frontal and left parietal clusters, which fell at the posterior end of the inferior frontal sulcus at its junction with the precentral sulcus (pIFS; MNI - 46, 0, 38) and in the intraparietal sulcus (IPS; MNI -38, -50, 44), respectively. The same target coordinates were used for all participants. For each participant, the two coordinates were transformed from MNI space into native anatomical space using the inverse deformation field from the unified segmentation of the T1 image within SPM12, and the resulting native-space coordinates were loaded into the neuronavigation software (Brainsight, Rogue Research Inc., Montreal, Canada) as stimulation targets. A third target, used only for determining the motor threshold, was placed manually on each participant’s T1 image over the “hand knob” of the left precentral gyrus (i.e., the hand area of motor cortex).

### Non-invasive brain stimulation

TMS was delivered with a MagStim Super Rapid^2^ Plus^1^ stimulator and a 70 mm figure-eight double air-film coil (MagStim, Whitland, United Kingdom). Coil position was tracked throughout all sessions with BrainSight v2.4.8 (Rogue Research Inc., Montréal, Canada) and a Polaris Spectra infrared camera (Northern Digital, Waterloo, Canada), after co-registering anatomical landmarks on the participant’s head to their T1 image. Participants wore earplugs during all stimulation.

In the first session we determined each participant’s resting motor threshold (MT). Surface electromyography (EMG) was recorded from the first dorsal interosseous muscle of the right hand. Starting from the hand-knob target, single pulses were delivered to the left primary motor cortex with the coil at ∼45° to the midline (handle pointing posterolaterally), and the coil was repositioned to locate the scalp site yielding the largest motor-evoked potential (MEP) (“motor hotspot”). MT was defined as the lowest stimulator output at which 5 of 10 consecutive pulses elicited an MEP of at least 70 µV peak-to-peak in the EMG trace or a visible twitch of the hand, with the muscle fully relaxed. Across the final sample, MT ranged from 54% to 85% of maximum stimulator output (*M*=67.3%, *SD*=9.2%).

In the two rTMS sessions, stimulation intensity was set to 110% of the individual MT for all participants. On every trial, a train of four biphasic pulses was delivered during the second half of the memory delay, beginning 1.0 s after the offset of the memory array plus a random delay drawn uniformly from 1–200 ms, so that the train never extended beyond 1.7 s into the 2 s delay (Figure 1B). Three stimulation patterns were used, intermixed across trials. In the theta condition the four pulses were delivered at 6 Hz (inter-pulse interval 166.7 ms; train duration 500 ms); in the alpha condition at 10 Hz (inter-pulse interval 100 ms; train duration 300 ms). In the arrhythmic control condition the first and last pulses were separated by 500 ms, matching the duration of the theta train, and the timing of the two intermediate pulses was drawn at random (1 ms resolution) on each trial, subject to the constraints that all three inter-pulse intervals lay between 20 and 400 ms and that the three intervals were not all within 20 ms of one another, which excludes quasi-rhythmic patterns. The arrhythmic condition thus delivered the same number of pulses within the same window of the delay as the rhythmic conditions, and served to control for non-frequency-specific effects of stimulation at each site (Riddle et al., 2020). Rhythmic trains were generated using the stimulator’s train mode, triggered from the task computer over a serial connection; arrhythmic trains were generated as individually timed single pulses triggered through a data acquisition device. Each session therefore comprised 960 pulses.

The stimulation trajectory was defined in BrainSight as perpendicular to the scalp at the target. The coil was held in place by an experimenter who monitored the coil position and corrected drifts in real time. The coil position relative to the planned trajectory was logged after every train. Trials in which the coil deviated by more than 5 mm from the target trajectory, in which one or more pulses were not delivered because of technical problems, or in which the trial was interrupted due to clear MEPs or twitches, eye twitching, or discomfort reported by the participant, were excluded from analysis. In total, 2.76% of trials were excluded (318 of 11,520; median 0.42% per session, *SD*=6.73%), leaving on average 233 trials per session (*SD*=16.2; range 150–240). Participants were monitored for discomfort throughout and were reminded that they could interrupt the session at any time.

## QUANTIFICATION AND STATISTICAL ANALYSIS

### Behavioral modeling

Response error was defined, for each of the six responses on a trial, as the angular distance on the color wheel between the reported and the target color (range -180 to 180). As a measure of accuracy we used the absolute error of the first response on each trial, averaged across trials within each condition. To separate the contributions of memory precision, binding errors (swaps), and guessing, we fitted the responses with the three-component mixture model of Bays and colleagues (Bays et al., 2009; Schneegans & Bays, 2016), implemented in the Analogue Report Toolbox, which describes the distribution of responses as a mixture of a von Mises distribution centered on the target color, von Mises distributions of the same concentration centered on the two non-target colors (swap errors), and a uniform distribution (random guesses). Parameters were estimated by maximum likelihood using expectation–maximization with multiple starting points. For each participant and session, the model was fitted separately to the trials of each condition, pooling all six responses per trial; the concentration parameter κ of the von Mises component was taken as the measure of memory precision.

To classify individual trials as swaps or guesses, we fitted the mixture model separately to the responses of each ordinal position (first to sixth) across all trials of a session and obtained, for each response, the posterior probabilities that it was generated by the target, non-target, or uniform component. A response was labelled a swap when its posterior probability of belonging to a non-target component exceeded 0.70, and a guess when its posterior probability of belonging to the uniform component exceeded 0.50. A trial was classified as a swap trial or a guess trial when at least three of its six responses received the corresponding label. The numbers of swap and guess trials were then counted within each condition. No trials were excluded from the analyses of precision or error based on these labels.

### Statistical analysis

To control for non-frequency-specific effects of stimulation within each site and session, and following the work by Riddle and colleagues (Riddle et al., 2020), all measures were expressed as difference scores between each rhythmic condition and the arrhythmic condition from the same session and presentation hemifield (theta minus arrhythmic; alpha minus arrhythmic). Each behavioral measure (κ, first-response absolute error, number of swap trials, number of guess trials) was submitted to a three-way repeated-measures analysis of variance (ANOVA) with the within-subject factors stimulation frequency (theta, alpha), stimulation site (pIFS, IPS), and presentation hemifield (left, right), implemented with the *fitrm* and *ranova* functions in MATLAB R2023b. Because the dependent variables were difference scores, the intercept term of each model tests whether rhythmic stimulation changed the measure relative to arrhythmic stimulation irrespective of frequency, site, or hemifield; main effects and interactions test whether this change depended on those factors. Significant effects were followed up with pairwise comparisons of estimated marginal means (*multcompare*), corrected for multiple comparisons using the Tukey–Kramer procedure. Effect sizes are reported as Cohen’s d, computed as the difference between conditions means divided by the pooled standard deviation (Cohen, 1992). The significant threshold was set at α=0.05 (two-tailed). All analyses were performed in MATLAB R2023b (The MathWorks Inc. Natick, USA).

## REFERENCES

Baddeley, A. D. (2012). Working Memory: Theories, Models, and Controversies. Annual Review of Psychology, 63(1), 1–29. 10.1146/annurev-psych-120710-100422

Baddeley, A. D., & Hitch, G. (1974). Working Memory. In Psychology of Learning and Motivation (Vol. 8, pp. 47–89). Elsevier. 10.1016/S0079-7421(08)60452-1

Barbosa, J., Babushkin, V., Temudo, A., Sreenivasan, K. K., & Compte, A. (2021). Across-Area Synchronization Supports Feature Integration in a Biophysical Network Model of Working Memory. Frontiers in Neural Circuits, 15, 716965. 10.3389/fncir.2021.716965

Bays, P. M., Catalao, R. F. G., & Husain, M. (2009). The precision of visual working memory is set by allocation of a shared resource. Journal of Vision, 9(10), 7. 1-11. 10.1167/9.10.7

Bender, A., Zhao, C., Vogel, E., Awh, E., & Voytek, B. (2025). Differential representations of spatial location by aperiodic and alpha oscillatory activity in working memory. Proceedings of the National Academy of Sciences, 122(30), e2506418122. 10.1073/pnas.2506418122

Beynel, L., Appelbaum, L. G., Luber, B., Crowell, C. A., Hilbig, S. A., Lim, W., Nguyen, D., Chrapliwy, N. A., Davis, S. W., Cabeza, R., Lisanby, S. H., & Deng, Z.-D. (2019). Effects of online repetitive transcranial magnetic stimulation (rTMS) on cognitive processing: A meta-analysis and recommendations for future studies. Neuroscience & Biobehavioral Reviews, 107, 47–58. 10.1016/j.neubiorev.2019.08.018

Bonnefond, M., & Jensen, O. (2012). Alpha Oscillations Serve to Protect Working Memory Maintenance against Anticipated Distracters. Current Biology, 22(20), 1969–1974. 10.1016/j.cub.2012.08.029

Christophel, T. B., Klink, P. C., Spitzer, B., Roelfsema, P. R., & Haynes, J.-D. (2017). The Distributed Nature of Working Memory. Trends in Cognitive Sciences, 21(2), 111–124. 10.1016/j.tics.2016.12.007

Cohen, J. (1992). A power primer. Psychological Bulletin, 112(1), 155–159. 10.1037/0033-2909.112.1.155

Dake, M., & Curtis, C. E. (2025). Perturbing human V1 degrades the fidelity of visual working memory. Nature Communications, 16(1), 2675. 10.1038/s41467-025-57882-8

D’Esposito, M. (2007). From cognitive to neural models of working memory. Philosophical Transactions of the Royal Society B: Biological Sciences, 362(1481), 761–772. 10.1098/rstb.2007.2086

D’Esposito, M., & Postle, B. R. (2015). The Cognitive Neuroscience of Working Memory. Annual Review of Psychology, 66(1), 115–142. 10.1146/annurev-psych-010814-015031

Foster, J. J., Sutterer, D. W., Serences, J. T., Vogel, E. K., & Awh, E. (2016). The topography of alpha-band activity tracks the content of spatial working memory. Journal of Neurophysiology, 115(1), 168–177. 10.1152/jn.00860.2015

Jabar, S. B., Sreenivasan, K. K., Lentzou, S., Kanabar, A., Brady, T. F., & Fougnie, D. (2023). Probabilistic and rich individual working memories revealed by a betting game. Scientific Reports, 13(1), Article 1. 10.1038/s41598-023-48242-x

Kiyonaga, A., Miller, J. A., & D’Esposito, M. (2024). Lateral prefrontal cortex controls interplay between working memory and actions (p. 2024.09.17.613601). bioRxiv. 10.1101/2024.09.17.613601

Kleiner, M., Brainard, D., & Pelli, D. (2007). What’s new in Psychtoolbox-3? https://pure.mpg.de/rest/items/item_1790332/component/file_3136265/content

Klimesch, W., Doppelmayr, M., Schwaiger, J., Auinger, P., & Winkler, T. H. (1999). ′Paradoxical’ alpha synchronization in a memory task. Cognitive Brain Research, 7(4), 493–501. 10.1016/S0926-6410(98)00056-1

Mackey, W. E., & Curtis, C. E. (2017). Distinct contributions by frontal and parietal cortices support working memory. Scientific Reports, 7(1), 6188. 10.1038/s41598-017-06293-x

Pagnotta, M. F., Santo-Angles, A., Temudo, A., Barbosa, J., Compte, A., D’Esposito, M., & Sreenivasan, K. K. (2026). Alpha phase coding supports feature binding during working memory maintenance. Communications Biology, 9(1), 922. 10.1038/s42003-026-10071-9

Pavlov, Y. G., & Kotchoubey, B. (2022). Oscillatory brain activity and maintenance of verbal and visual working memory: A systematic review. Psychophysiology, 59(5), e13735. 10.1111/psyp.13735

Preston, M., Schaworonkow, N., & Voytek, B. (2025). Time-Resolved Aperiodic and Oscillatory Dynamics during Human Visual Memory Encoding. Journal of Neuroscience, 45(16). 10.1523/JNEUROSCI.2404-24.2025

Riddle, J., Scimeca, J. M., Cellier, D., Dhanani, S., & D’Esposito, M. (2020). Causal Evidence for a Role of Theta and Alpha Oscillations in the Control of Working Memory. Current Biology, 30(9), 1748–1754.e4. 10.1016/j.cub.2020.02.065

Rossi, S., Hallett, M., Rossini, P. M., & Pascual-Leone, A. (2009). Safety, ethical considerations, and application guidelines for the use of transcranial magnetic stimulation in clinical practice and research. Clinical Neurophysiology, 120(12), 2008–2039. 10.1016/j.clinph.2009.08.016

Schneegans, S., & Bays, P. M. (2016). No fixed item limit in visuospatial working memory. Cortex, 83, 181–193. 10.1016/j.cortex.2016.07.021

van Engen, Q., Chau, G., Smith, A., Adam, K., Donoghue, T., & Voytek, B. (2026). Dissociating Contributions of Theta and Alpha Oscillations from Aperiodic Neural Activity in Human Visual Working Memory. Journal of Neuroscience, 46(7). 10.1523/JNEUROSCI.2340-24.2025

Wallis, G., Stokes, M., Cousijn, H., Woolrich, M., & Nobre, A. C. (2015). Frontoparietal and Cingulo-opercular Networks Play Dissociable Roles in Control of Working Memory. Journal of Cognitive Neuroscience, 27(10), 2019–2034. 10.1162/jocn_a_00838

Wianda, E., & Ross, B. (2019). The roles of alpha oscillation in working memory retention. Brain and Behavior, 9(4), e01263. 10.1002/brb3.1263

Yarkoni, T., Poldrack, R. A., Nichols, T. E., Van Essen, D. C., & Wager, T. D. (2011). Large-scale automated synthesis of human functional neuroimaging data. Nature Methods, 8(8), 665–670. 10.1038/nmeth.1635

